# Interneuron-like features of microglia in the mouse outer retina

**DOI:** 10.1101/2024.01.12.575392

**Authors:** Sara Toros, Thomas Kane, Aura Hare, Hassina Abbar, Akshay Madoo, Giulia De Rossi, Kate Powell, Matteo Rizzi

## Abstract

Microglia have recently been associated with complex functionalities both in the brain and the retina, including in synaptic function and plasticity. While investigating the morphology, protein expression profile and connectivity of microglia in the mouse outer retina, we have found that they closely resemble interneurons in the outer retina of other species. We argue that functional studies are needed to reveal whether they also perform an interneuron-like role in vision.

## Introduction

Glia, first described by Virchow as passive neural elements and the ‘connective tissue’ or ‘glue’ of the brain^1^, were long considered as inert bystanders in the central nervous system (CNS) formation and functionality. It is now widely accepted that glial functions are far more complex than initially described (reviewed in^2,3^). For example, recent evidence suggests that a subtype of astrocytes selectively expresses synaptic-like glutamate-release machinery, which enables Ca^2+^ dependent exocytosis, a feature shared with neurons^4^. Similarly, microglia have been shown to play a more complex role in the brain and the retina, which goes beyond the historical perception as a macrophage-like cell. Among many functions, they display distinct activation states in homeostasis and disease^5^, have an important function in synapse remodelling during development and plasticity^6-10^, and significantly contribute to synapsis maintenance and the integrity of the mature retina^11^.

In the retina, microglia primarily reside in two distinct synaptic layers, the outer and inner plexiform layers (OPL and IPL, respectively)^12^. In the OPL of cats and primates, microglia morphology has been described as surprisingly similar to that of axon-less H2 horizontal cells, extending processes towards photoreceptors^13^. In the mouse retina, microglia exhibit spatial variation in their dependency on interleukin-34 (IL-34), and this differential feature has been exploited to isolate and dissect their distinct functional roles^5^. IL34-mediated IPL microglia depletion was shown to specifically reduce cone-driven bipolar cell activation^5^, suggesting that IPL microglia might perform a pathway-specific modulation within the retinal circuitry^5^. Broader depletion of all retinal microglia led to a reduction in both a- and b-wave responses^11^, indicating that OPL microglia might play a specific role in modulating photoreceptor function. Taken together, these studies suggest that distinct sub-populations of microglia might contribute to specific aspects of visual processing.

Given the similarity in morphology and function between microglia and some types of interneurons, we aimed to characterise the expression profile, connectivity, and spatial distribution of microglia in the mouse outer retina.

## Results

We first performed an immunohistochemical characterisation, and as previously reported^14,15^, we confirmed that OPL microglia express the microglial marker Iba1 (Fig.1a). We then looked at classic interneuron markers and found that OPL microglia expressed both Calretinin and Parvalbumin (Fig.1b), which are markers of central nervous system interneurons^16^, including retinal amacrine cells^17,18^ and horizontal cells^19,20^. We found that OPL microglia did not express Calbindin, which selectively identifies the H1 subtype of horizontal cells in mouse^21^. However, the OPL microglia pool did exhibit expression of Isl1 (Fig.1c), a horizontal cell marker used to identify the H2 horizontal cell subtype in other species^22-25^. This is an interesting finding, because H2 cells have not, so far, been identified in mouse^26^. These results suggest that, in terms of protein markers, microglia in the mouse OPL have a profile consistent with cells that have been identified as H2 cells in other species, including human, and differ in expression profile from mouse H1 cells (Fig.1d).

**Figure 1.**
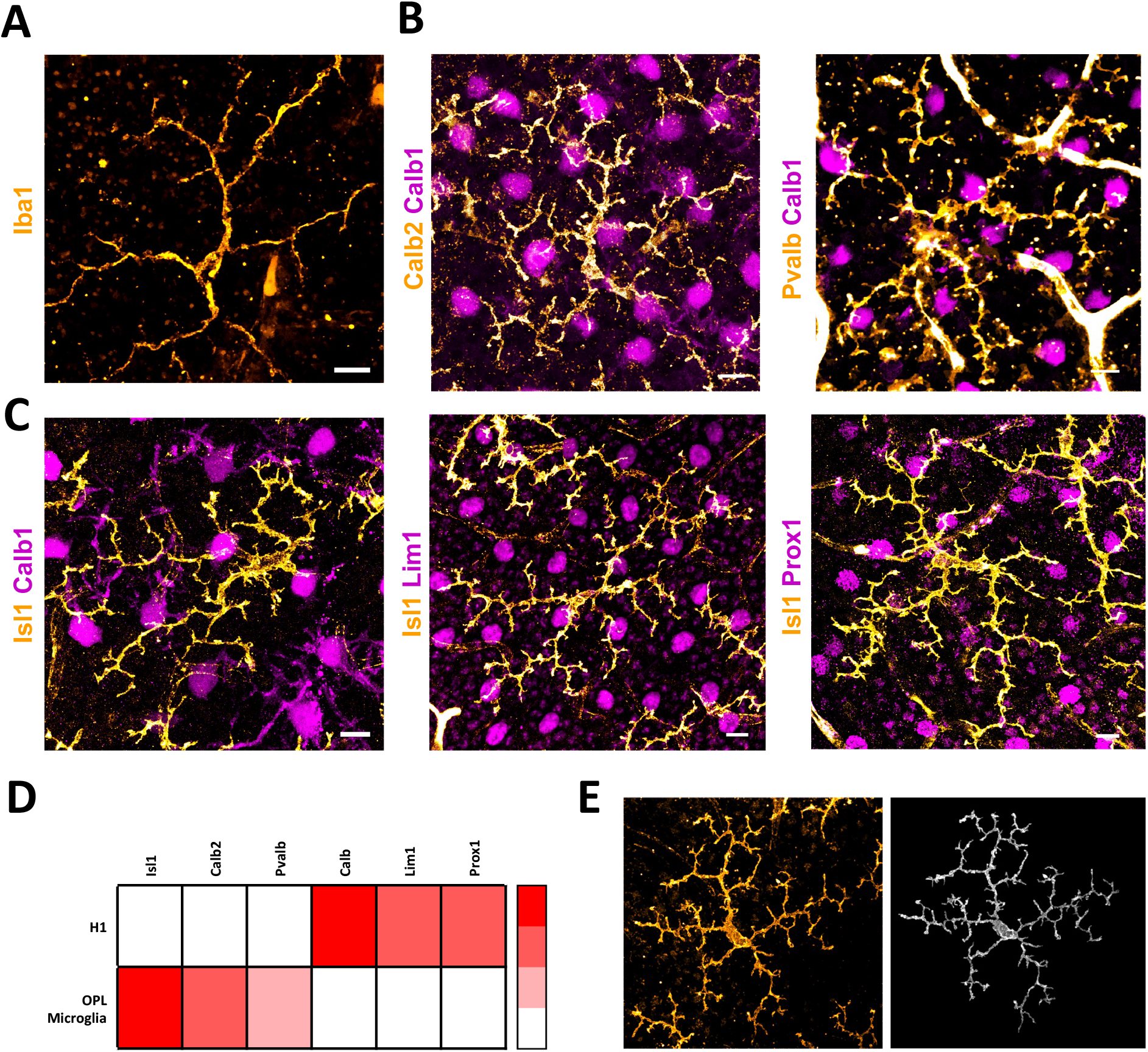
Microglial cells in the OPL express common interneuron markers. **A)** Microglia in the outer plexiform layer (OPL) express the common microglial marker Iba1. **B, C)** Flat mounted mouse retinas imaged at the OPL level. Maximum intensity projections showing the selective expression of interneuron markers in H1 cells (magenta) and microglia (orange). Note that the images are maximum projections of two different depths to show cell bodies of both cell types. **D)** Relative expression of six common interneuron markers in H1 cells and microglia reveal differential profiles in the two cell types. **E)** Immunodetection of a microglial cell in the OPL (Isl1 staining) and corresponding 3D reconstruction. Scale bars: 10 μm.

Laterally projecting horizontal cells lie horizontally in the OPL and provide feedback inhibition to photoreceptors. Of the two horizontal cell subtypes, H1 cells send axon terminals to rod synapses, while H2 cells make contacts with cone pedicles^27^. Likewise, OPL microglia lie horizontally within the synaptic layer, spanning a thickness of only ∼10 μm. 3D reconstructions of the soma and lateral processes (Fig.1e) show that they extend horizontally, except for calyx-like structures elongating towards photoreceptors (Fig.2). Interestingly, microglia processes appeared to terminate in the layer containing cone terminals, not extending as far as rod photoreceptor pedicles (Fig.2a). We analysed their connectivity features by labelling Isl1 in microglia in both 6 weeks old Chrnb4.EGFP mice, a transgenic mouse line with GFP^+^ cones (Fig.2a, b), and in wild type mice, co-staining the retina targeting the cone marker cone arrestin (Fig.2c). We found that microglial dendritic processes extend towards the cone pedicles, revealing putative sites of contact (Fig.2). We generated 3D surface renderings (Imaris software, see Methods) of cone pedicles and microglial processes (Fig.2d, e). Analysis of the surface contact area between cone pedicles and microglia unveiled regions of contact between the two (average surface contact 2.06±2.27 μm^2^), on the lateral surface of the terminals, outside of the triad synapse (2d,e). Notably, H2 cells in other species have also been shown to contact cone pedicles on their lateral aspect^28^.

**Figure 2.**
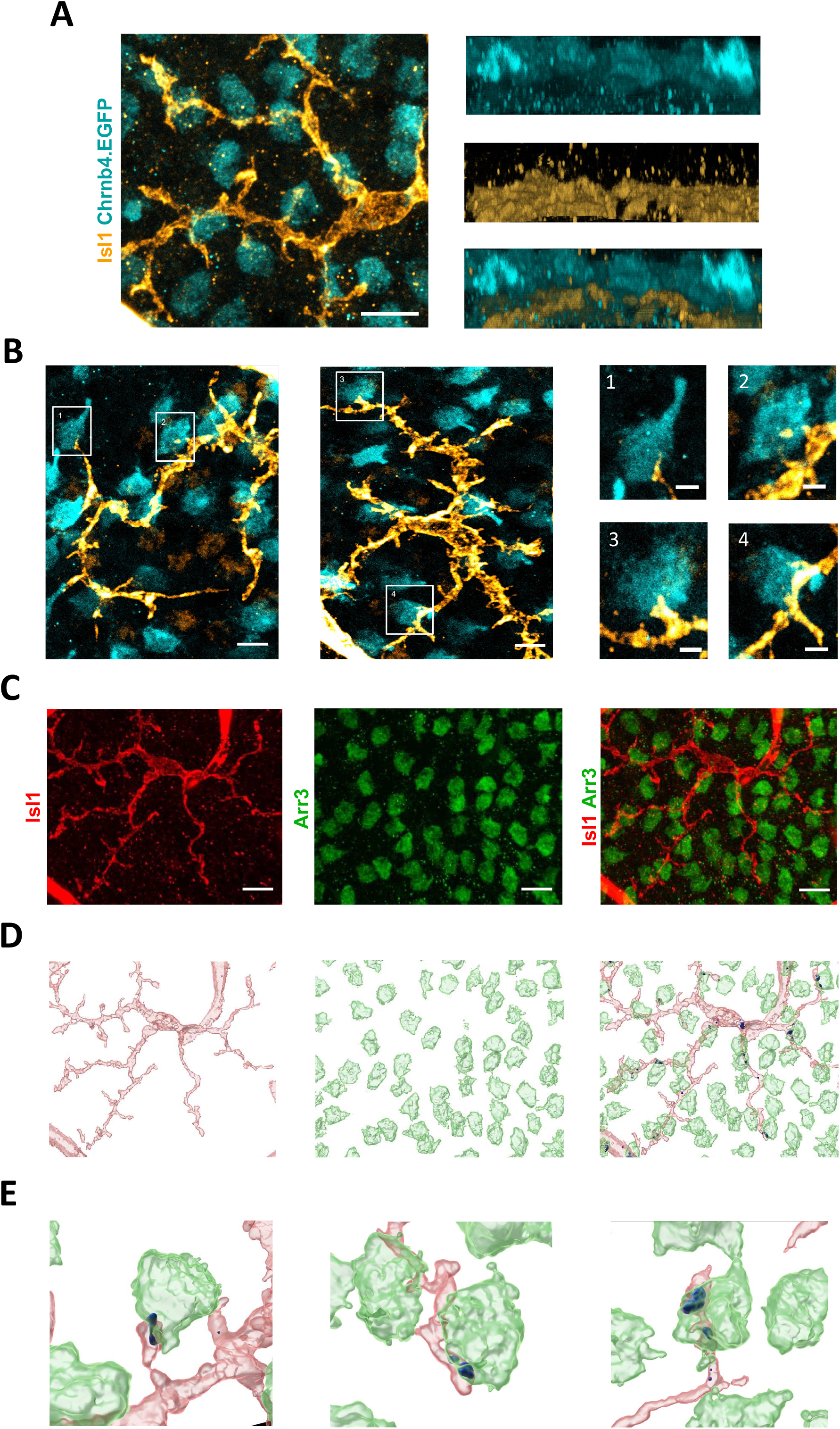
Microglial cells connect laterally to cone terminals in the outer plexiform layer. **A)** Maximum projection and lateral perspective of the outer plexiform layer (OPL) illustrating the position of microglia in relation to the cone terminals. Processes were found to reach cone terminals but not to extend further towards rod terminals. **B)** Low and high magnification of maximum intensity projection showing microglial processes contacting cone pedicles. **C)** Maximum intensity projection showing a microglial cell in the OPL and cone terminals. **D, E)** Low and high magnification respectively of the 3D surface rendering of the microglia cell, cone terminals and their contacts (see Methods). Note the surface contact area between the cone terminals and microglial lateral processes (blue). Microglia connect to the cone terminals outside of their synaptic cleft and on the lateral surface, much like H2 cells^28^. Scale bars: A, B, C, D: 10 μm, B1, B2, B3, B4: 0.3 μm

We then analysed the OPL microglia at a population level, focusing on cellular patterning and distribution (Fig.3). To assess the regularity of microglia in the OPL, we compared our real distribution to a random generated cell simulation, density-matched and constrained for cell soma size (Fig.3b). The nearest neighbour distances (NN) and Voronoi domain areas (VD) analyses, with their regularity indexes (NNRI and VDRI, respectively) (Fig.3c), are spatial statistical parameters commonly used to describe the spatial regularity of a cell population^29^. Both metrics (NN and VD) and their relative regularity indexes show that the spatial distribution of OPL microglia differs from random, suggesting a regular mosaic as their patterning model (Fig.3c).

**Figure 3.**
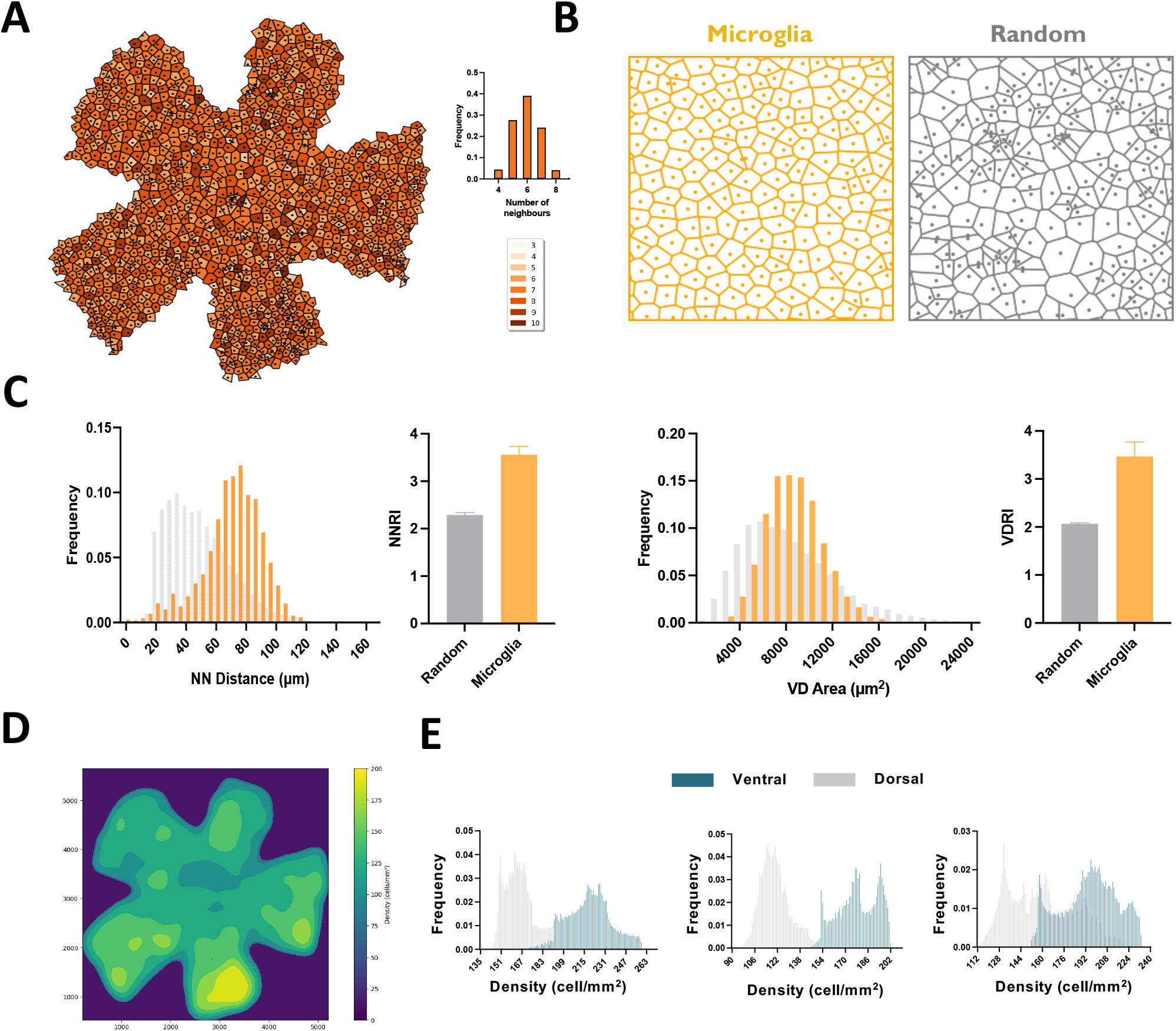
Spatial statistical analyses of OPL microglia patterning and distribution. **A)** Voronoi domain (VD) areas of OPL microglia across a mouse retina, colour-coded for number of neighbours. Bar plot indicates frequency of number of neighbours. **B)** Real versus simulated soma arrays showing VD areas, in orange and grey, respectively. Distributions are matched for number of cells, density and soma size. **C)** Frequency distributions of VD and nearest neighbour (NN) and their respective regularity indexes, comparing microglia distribution from our data and random distributions from computer simulations. **D)** Representative density plot of OPL microglia for the same retina shown in A, which follow a dorsal-ventral gradient in the mouse retina (the retina is oriented such that dorsal is at the top of the image; orientation was based on the S opsin gradient). **E)** Frequency distributions of OPL microglia densities in dorsal-most (grey) and ventral-most (blue) regions of three mouse retinas.

We used the same two-dimensional point patterns to analyse the density of OPL microglia in the mouse retina (Fig.3d). We found a dorsal-ventral gradient, with peak density of OPL microglia in the central ventral retina (Fig.3d,e). Interestingly, the S-cone pathway has been shown to follow a very similar dorsal-ventral gradient in the mouse retina, where both S-cones and S-cone bipolars peak in density in the ventral-most part of the retina^30^. These results suggest that OPL microglia follow a pattern that closely matches that of S-cones.

## Discussion

Recent evidence suggests that glial cells might possess some key features that were typically thought of as neuron-specific^6-10^, including their ability to affect synaptic transmission^5,11^. In the mouse retina, the IPL microglia pool has been shown to play a functional role in processing visual stimuli, suggesting a possible interneuron-like function of these cells^5^. Even though the OPL of the vertebrate retina is known to have multiple horizontal neurons in many species^25^, only one has been characterised in the mouse^26^. While investigating the expression profile of microglia in the mouse OPL, we surprisingly found that the microglia of this synaptic layer express common interneuron markers, which are associated with horizontal cells in other species. In particular, the Isl1 marker has been used to identify H2 horizontal cells in species such as chicken and human^22-24^.

Our 3D analysis of mouse microglia and cone connections showed their calyceal processes contacting cone terminals laterally, similarly to what has been shown for H2 cells in non-human primates, where the lateral connections formed by H2 cells are distinct from H1 connections with the triad synapse^28^.

In many species, including non-human primates and humans, H2 cells are mainly believed to be dedicated to compute visual stimuli originating from S cones^31,32^. Notably, our study revealed that OPL microglia follow a gradient similar to that of S-cones^30^, in which both cell types are highly concentrated in the ventral retina. This pattern suggests the potential involvement of OPL microglia in similar lateral interactions. A microglia dorsal-ventral gradient has not been observed in other glial populations, which have instead been shown to have a homogeneous density throughout the retina^33^.

Interestingly, the morphology and phenotype of the OPL mouse microglia closely resemble the A-type horizontal cell found in other species. In fact, as previously reported, microglia in other species have been confused with axon-less horizontal cells, both for morphological similarity and connectivity features to cones^13^. Therefore, it might be interesting to re-evaluate the distinction between OPL microglia and H2 axon-less cells by co-staining with markers used to profile both cell types.

In this study, we report OPL microglia features that closely resemble those of H2 cells in other species. Our findings, taken together with other recent discoveries^4,5,11^, suggest that the clear demarcation between the concepts of ‘neurons’ and ‘glia’ should be revisited. Complementing this morphological and structural study with functional tests will allow for a better characterisation of the role of microglia in processing visual stimuli in the retinal pathways.

## Materials & Methods

### Animal handling and ethic statement

All experiments in mouse were approved by the local Animal Welfare and Ethics Review Board (UCL, London, UK) and the Home Office and conformed to the guidelines on the care and use of animals adopted by the Society for Neuroscience and the Association for Research in Vision and Ophthalmology (Rockville, MD, USA).

### Tissue collection and immunohistochemistry

Eyes were rapidly enucleated, and immersion fixed in 4% paraformaldehyde (PFA, Thermo Scientific) overnight at 4°C. Retinas were dissected free of the eye and incubated with primary antibodies in PBS with 3% Triton X-100 (Sigma Aldrich) overnight at 4°C. Retinas were washed 3-4 times with PBS and incubated with secondary antibodies in the permeabilization solution, overnight at 4°C. Retinas were thoroughly washed with PBS and counterstained with DAPI (Thermo Fisher, D1306) before being immersed in a clearing solution previously used (Cristante et al., 2018) either overnight at 4°C or 2h at room temperature. Retinas were then flat-mounted by making 4 cuts from the retinal periphery to the optic nerve, and coverslipped with the same clearing solution as mounting medium.

Primary antibodies used were (dilution 1:200): goat anti-Iba1 (Abcam, ab5076), rabbit anti-Calbindin (Abcam, ab108404), mouse anti-calretinin (Sigma, MAB1568), mouse anti-parvalbumin (Sigma, P3088), mouse anti-Isl1 (DSHB, 40.2D6-c), rabbit anti-Lhx1 (Abcam, ab229474), rabbit anti-Prox1 (Sigma, AB5475), rabbit anti-Arr3 (Abcam, ab15282), goat anti-S opsin (Rockland, 600-101-MP7).

Secondary antibodies used were (dilution 1:1000, Thermo Fisher Scientific): donkey anti-mouse Alexa-Fluor 647 (A-31571), goat anti-rabbit Alexa-Fluor 647 (A-21245) and 594 (A32740), donkey anti-goat Alexa-Fluor 488 (A32814).

### Microscopy and Image analysis

Images were acquired on a Leica STELLARIS 8 confocal microscope equipped with a computer-driven motorised stage, controlled by the LAS X acquisition and analysis software (Leica Microsystems, Wetzlar, Germany). Both high-resolution z-stacks and automatically merged tile scans of whole retinas were acquired with a 40x oil objective, the latter using the LAS X-integrated Navigator tool.

3D reconstructions of single microglial cells were obtained semi-automatically using the SNT plugin in Fiji/ImageJ 2.9.0.

To analyse the contact areas between microglia and cone pedicles, z-stacks confocal images were 3D rendered using Imaris 10.1.1 (Bitplane, South Windsor, CT, USA). X and Y signals were used to create Imaris surfaces for microglia and cone pedicles, respectively. The overlapping area between the two surfaces was masked and shown as an additional channel. The contact areas between microglia and cone pedicles were analysed and the cumulative surface area was automatically calculated by the software.

To analyse the cell patterning and distribution of the OPL microglia pool, OPL Isl1^+^ microglia were manually quantified in automatically merged tile scans of n = 3 retinal whole mounts using the multi-point tool in Fiji/ImageJ 2.9.0. X and Y coordinates of each OPL Isl1^+^ cell were extracted, and soma patterns were generated. NN distances and VD areas were calculated for each retinal sample, and their regularity indexes were computed as the ratio of their mean to their standard deviation. To minimise edge effects on the computation of these spatial statistics, the values of the cells closer to the outer edge of the whole mount were discarded. To acquire accurate statistics, but also allow analysis of the maximum number of cells by removing the minimum number of edge cases, the concept of ‘alpha shapes’, a generalisation of the concept of the convex hull, was used. This shape was defined using a value for α that is equal to the average inter-cell distance (ICD) of the distribution, generating a border to which a morphological dilation using a value of the ICD/2 was applied. Excluding only cells with a Voronoi domain vertex that lies outside of this region gave us the correct population of cells for the analysis.

Random cell distributions were produced for statistical and visual comparison with the microglia population. The same density was used, and a minimum cell separation equal to the size of the microglia cell body. The statistics for the random distributions are averages of hundreds of iterations of this random simulation.

To analyse the density of the OPL microglia pool, the same soma two-dimensional point patterns were analysed in Python and density plots were generated. We used gaussian kernel density estimation to acquire a probability density function for microglia over the whole retina. We then selected the most ventral and most dorsal regions according to the S opsin gradient and analysed the density distribution in those areas.

Voronoi diagrams, alpha shapes, random simulations, and all other computations were generated using our own Python modules. A range of libraries were used in creating these, including scipy, shapely, imantics, and network.

## Acknowledgments

We would like to thank Jennifer Sun for eyes from PFA-perfused mice and members of our lab for helpful discussion and for critically reading the manuscript. Work was supported by funding to M Rizzi and K Powell by Moorfields Eye Charity, Fight for Sight, UCL Tech Fund.

## Conflict of interests

all authors declare no conflict of interests.

## Notes

### Competing Interest Statement

The authors have declared no competing interest.

